# Stability Oracle: A Structure-Based Graph-Transformer for Identifying Stabilizing Mutations

**DOI:** 10.1101/2023.05.15.540857

**Authors:** Daniel J. Diaz, Chengyue Gong, Jeffrey Ouyang-Zhang, James M. Loy, Jordan Wells, David Yang, Andrew D. Ellington, Alex Dimakis, Adam R. Klivans

## Abstract

Stabilizing proteins is a fundamental challenge in protein engineering and is almost always a prerequisite for the development of industrial and pharmaceutical biotechnologies. Here we present Stability Oracle: a structure-based graph-transformer framework that achieves state-of-the-art performance on predicting the effect of a point mutation on a protein’s thermodynamic stability (ΔΔG). A strength of our model is its ability to identify *stabilizing* mutations, which often make up a small fraction of a protein’s mutational landscape. Our framework introduces several data and machine learning innovations to overcome well-known challenges in data scarcity and bias, generalization, and computation time. Stability Oracle is first pretrained on over 2M masked microenvironments and then fine-tuned using a novel data augmentation technique, Thermodynamic Permutations (TP), applied to a ∼120K curated subset of the mega-scale cDNA display proteolysis dataset. This technique increases the original 120K mutations to over 2M thermodynamically valid ΔΔG measurements to generate the first structure training set that samples and balances all 380 mutation types. By using the masked microenvironment paradigm, Stability Oracle does not require a second mutant structure and instead uses amino acid structural embeddings to represent a mutation. This architectural design accelerates training and inference times: we can both train on 2M instances with just 119 structures and generate deep mutational scan (DMS) predictions from only the wildtype structure. We benchmark Stability Oracle with both experimental and AlphaFold structures of all proteins on T2837, a test set that aggregates the common test sets (SSym, S669, p53, and Myoglobin) with all additional experimental data from proteins with over a 30% sequence similarity overlap. We used TP augmented T2837 to evaluate performance for engineering protein stability: Stability Oracle correctly identifies 48% of stabilizing mutations (ΔΔG < −0.5 kcal/mol) and 74% of its stabilizing predictions are indeed stabilizing (18% and 8% of predictions were neutral and destabilizing, respectively). For a fair comparison between sequence and structure-based fine-tuned deep learning models, we build on the Prostata framework and fine-tune the sequence embeddings of ESM2 on our training set (Prostata-IFML). A head-to-head comparison demonstrates that Stability Oracle outperforms Prostata-IFML on regression and classification even though the model is 548 times smaller and is pretrained with 4000 times fewer proteins, highlighting the advantages of learning from structures.

## 1 Introduction

The ability to predict and understand the change in protein thermodynamic stability (ΔΔ*G*) for an amino acid substitution is a core task for the development of protein-based biotechnology, such as industrial biocatalysts [1, 2, 3] and pharmaceutical biologics [4, 5, 6, 7]. Proteins with enhanced thermodynamic stability are less prone to unfolding and aggregation and are more engineerable [8, 9]; stabilizing the scaffold of a protein enables downstream exploration of potentially destabilizing mutations that may improve a target function [8]. Thermodynamic stability is measured by the change in Gibbs free energy (Δ*G*) between the native and unfolded state and reflects the underlying integrity of the global structure. Engineering proteins is a very laborious process and bottlenecks the development of protein-based biotechnologies[10, 11], making the development of computational methods that can accurately predict ΔΔ*G* of a point mutation, and in turn identify stabilizing mutations, a highly active research area[12, 13, 14, 15, 16].

Deep learning is currently revolutionizing many physical and biological disciplines [17, 18, 19, 20, 21], with AlphaFoldV2 leading the way as the scientific breakthrough of the year in 2021 [22, 23] and sparking an entire wave of deep learning structure prediction tools [24, 25]. Although a variety of sequence-based [26, 27] and structure-based [28, 29, 30] deep learning frameworks for stability prediction have been reported, the lack of data and machine learning engineering issues have prevented deep learning algorithms from having a similarly revolutionary impact on protein stability prediction (See Appendix C for details on the current deep learning frameworks).

Systematic analyses [15, 13, 16, 31] of the state-of-the-art computational stability predictors published over the last 15 years highlight data scarcity, variation, bias, and leakage, and poor metrics for evaluating model performance as the critical issues hindering meaningful progress (See Appendix D for summary on the primary data issues). As a result, the community still primarily relies on knowledge-based methods, such as Rosetta [32] and FoldX [33], or shallow machine learning approaches [34, 35, 36, 37, 38, 39, 40, 41, 42]. Furthermore, third-party analyses emphasize that only about 20% of predicted stabilizing mutations from the current state-of-the-art computational tools are actually stabilizing, even though 75-80% accuracy is often reported upon publication [14, 31, 13]. This poor generalization to new datasets highlights how pervasive the data leakage between train-test splits is in the computational stability prediction community [16, 31, 14, 13]. Finally, current computational tools are evaluated with Pearson correlation and RMSE as the primary metrics. Due to the significant imbalance between stabilizing (<30%) and destabilizing mutations in training and test sets and the innate variations associated with measuring ΔΔ*G* [15], improvements in these metrics do not necessarily translate into improvements for identifying stabilizing mutations [31, 13]. Thus, metrics such as precision, recall, area under receiver operating characteristic (AUROC), and Matthew’s correlation coefficient (MCC) should be prioritized to monitor improvements for protein engineering applications [31, 13, 14].(See Appendix D for a summary on metric issues.)

To address these issues and develop a robust stability prediction model for protein engineering, we present Stability Oracle: a structure-based deep learning framework that makes use of several innovations in data and machine learning engineering specific to stability prediction. Stability Oracle uses a graph-transformer architecture that treats atoms as tokens and utilizes their pairwise distances to inject a structural inductive bias into the attention mechanism. The input to Stability Oracle consists of the local chemistry surrounding a residue with the residue deleted (masked microenvironment) and two amino acid embeddings to represent a specific point mutation. This design decision enables Stability Oracle to generate all 380 possible point mutations from a single microenvironment, circumventing the need for computationally generated mutant structures and making deep mutational scanning inference computationally inexpensive.

To improve the generalization capability of Stability Oracle, we introduce a data augmentation technique—Thermodynamic Permutations (TP). Thermodynamic Permutations, similar to Thermodynamic Reversibility (TR)[43], are based on the state function property of Gibbs free energy [44]. For a specific position in a protein, TP expands *n* empirical ΔΔ*G* measurements into *n*(*n −* 1) thermodynamically valid measurements, thus increasing the dataset by up to an order of magnitude based on the number of residues that have multiple amino acids experimentally characterized (Figure 11, Appendix A).

TP enables all 380 mutation types to be sampled in a training and test set for the first time, alleviating common data biases such as those produced by alanine scanning experiments. Furthermore, unlike TR, it generates a balanced set of stabilizing and destabilizing mutations to non-wildtype amino acids. This allows us to better assess generalization for protein engineering, as the goal is to mutate away from wildtype.

We present three new datasets (C2878, cDNA120K, T2837) to address data leakage issues. MMseqs2 [45] was used to generate the three datasets and ensure the maximum overlap between proteins in the training and test set is below 30% sequence similarity. Concat 2878 (C2878) and Test 2837 (T2837) datasets are new training and test splits from previously established training and test sets, respectively. The third (cDNA120K) is a curated subset of the cDNA display proteolysis dataset #1[46], a dataset of *∼*850K thermodynamic folding stability measurements across 354 natural and 188 de novo mini-protein domains (40-72 amino acids). This latter dataset is especially interesting because of its size and because it relies upon proteolytic stability as a surrogate for thermodynamic stability.

Finally, we generate a sequence-based counterpart for Stability Oracle by fine-tuning ESM2 on our curated training and test sets using the Prostata framework [26] and train Prostata-IFML. With Prostata-IFML, we conduct head-to-head comparisons to demonstrate the advantage of structure over sequence-based methods. Overall we show that Stability Oracle and Prostata-IFML are state-of-the-art structure and sequence-based frameworks for computational stability prediction, respectively.

## 2 Results

### 2.1 Designing a Graph-Transformer Framework for Structure-based Protein Engineering

In prior work, we have experimentally shown that representations learned via self-supervised deep learning models on masked microenvironments (MutCompute and MutComputeX) can identify amino acid residues incongruent with their surrounding chemistry. These models can be used to “zero-shot” predict gain-of-function point mutations [47, 48, 49], including in protein active sites of computational structures [50]. Self-supervised models, however, generate mutational designs that are not biased towards a particular phenotype and do not update predictions based on experimental data [19].

The MutCompute framework uses a voxelized molecular representation [47, 50]. For protein structures, voxelization is a suboptimal molecular representation, as most voxels consist of empty space, and rotational invariance property is not encoded. The MutCompute frameworks use convolution-based architectures, which lag behind modern attention-based architectures in terms of representation learning and predictive power.

To develop a more powerful and generalizable framework for downstream tasks, we first built MutComputeXGT, a graph-transformer version of MutComputeX (Figure 1a). Each atom is represented as a node with atomic elements, partial charges, and SASA values as features with pairwise atomic distances labeling edges. Our graph-transformer architecture converts the pairwise distances into continuous and categorical attention-biases to provide a structure-based inductive bias for the attention mechanism. To generate likelihoods for the masked amino acid, we average the embeddings of all atomic tokens within 8Å of the masked C*_α_*. The design decision to narrow the pooling to atoms in the first contact shell of the masked amino acid is based on insights from systematic variation of the microenvironment volume when training self-supervised 3DCNNs [51]. With a similar number of parameters and the same train-test splits, MutComputeXGT demonstrates superior representation learning capacity than MutComputeX, achieving wildtype accuracy of 92.98%*±*0.26% compared to *∼*85% [50].

**Figure 1:**
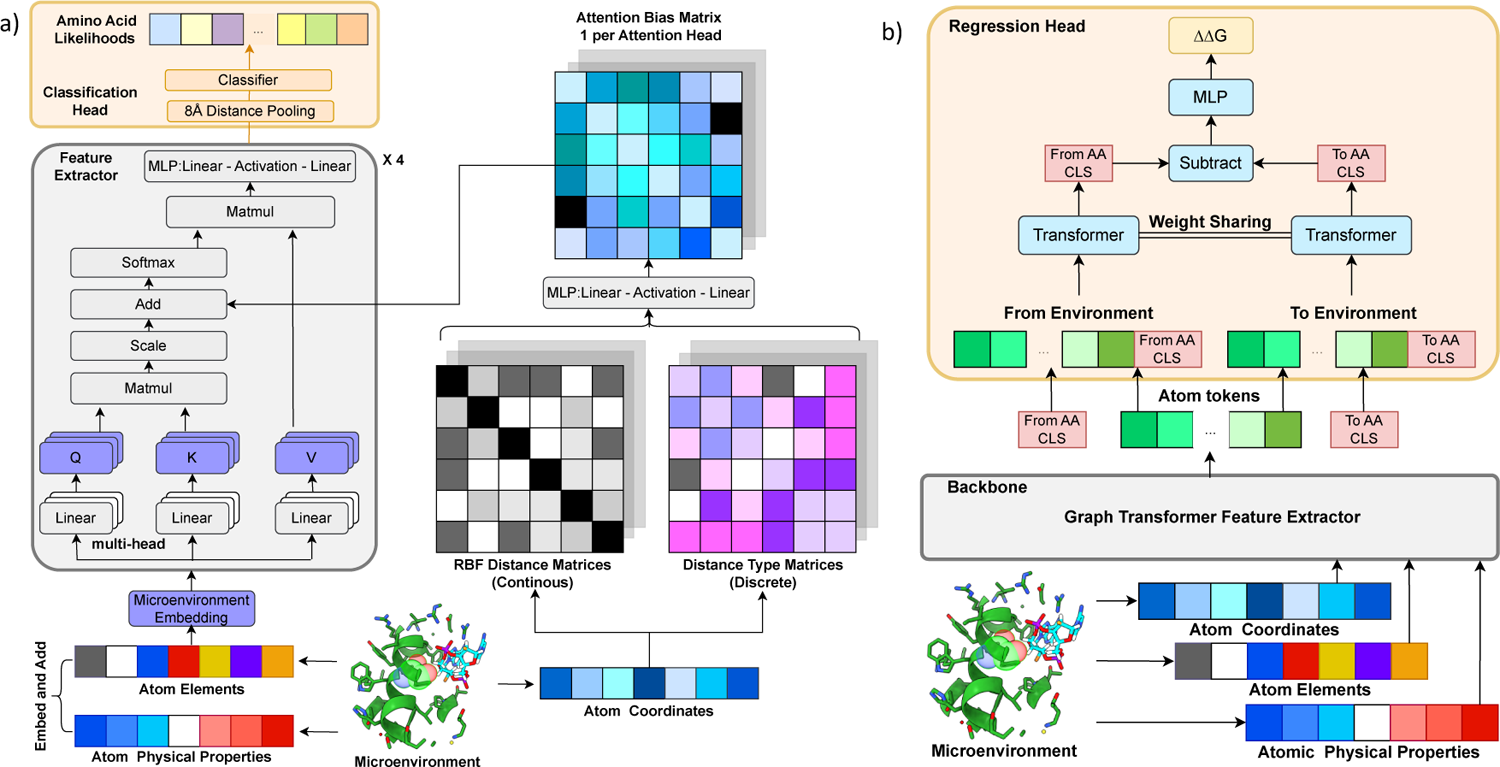
Overview of model architectures. a) Self-supervised pre-training graph-transformer architecture (MutComputeXGT). b) Fine-tuning pre-trained graph transformer for stability regression (Stability Oracle). The “from” AA and “to” AA CLS tokens are the corresponding weights of the last linear layer in the classifier of MutComputeXGT for that amino acid.

The Stability Oracle architecture makes use of both the feature-extractor and classifier of MutComputeXGT for supervised fine-tuning (Figure 1b). Previous structure-based stability predictors [28, 29] require two structures, one for the wildtype and one for the mutant. Instead, Stability Oracle uses only one structure encoded via a MutComputeXGT representation, and two amino acid level embeddings, one for the “from” and one for the “to” amino acid. This architectural decision alleviates the computational burden of generating a mutant structure for each ΔΔ*G* prediction: for a typical 300 amino acid protein, prior work would generate 5700 computational structures (from Rosetta [32] or AlphaFold [22]) in order to predict the ΔΔ*G* of every possible single point mutation. Stability Oracle, on the other hand, requires only one structure and can predict the ΔΔ*G* of all 20 amino acids at a specific residue in *∼*50 ms. Performance metrics on proteins of varying length are provided in Table 1, Appendix A.

The “from” and “to” amino acid embeddings are the weights from the final layer of the MutComputeXGT classifier. This design decision is based on the insight that the weights of these 20 neurons represent the similarity of a microenvironment’s features to each of the 20 amino acids prior to being normalized into a likelihood distribution. Thus, they are structure-based contextualized embeddings of the 20 amino acids learned from a 50% sequence similarity representation of the Protein Data Bank (PDB)[52], we will refer to them as structural amino acid embeddings. We observe that structural amino acid embeddings significantly improve performance compared to the naïve one-hot representation (Table 2, Appendix A).

We highlight several design decisions of the regression head used in Stability Oracle. Of note is the use a Siamese attention architecture that treats the mutation embeddings as two classification (CLS) tokens (Figure 1b). CLS tokens are commonly used in the natural language processing (NLP) community to capture the global context of the word-level token inputs for downstream tasks [53]. Since atoms and amino acids are chemical entities at different scales, we designed the regression head to contextualize the “from” and “to” amino acid-level CLS tokens with the atomic tokens of a particular microenvironment. Once contextualized for a given microenvironment, the two amino acid CLS tokens are subtracted from each other and decoded into a ΔΔ*G* prediction. We chose to subtract the two amino acid CLS tokens to enforce the state function property of Gibbs free energy [54], providing the proper inductive bias for Thermodynamic Reversibility and “self-mutations” (where ΔΔ*G* = 0 kcal/mol).

### 2.2 Training Stability Oracle to Generalize Across Protein Chemistry

The Stability Oracle framework was designed to generalize across all 380 mutation types at all positions within a protein structure. The development of such a model has historically been limited by data scarcity, bias, and leakage. To address these issues, we curated new training and test datasets using a new data augmentation technique called Thermodynamic Permutations (TP).

It is well known that a major issue with prior works is the inclusion of similar proteins in both the training and test set (“data leakage”), resulting in poor generalization. It has been demonstrated that train-test splits at the mutation, residue, or protein level result in overfitting and strict sequence clustering is required to mitigate this effect [14, 55, 13, 31]. Here we created new train-test splits based on a 30% sequence similarity threshold computed by MMSeqs2 [45]. First, we designed T2837: a test set that combines the common test sets (SSym [54], S669[12], Myoglobin, and P53) with any additional experimental data in common training sets (q1744[28], o2567 [16], s2648[38], and FireProtDB [56]) with a sequence similarity greater than 30%. After establishing the T2837 test set, we generate our training set (C2878) from the remaining experimental data by filtering out proteins with over 30% sequence similarity to T2837. The same procedure was used to construct the cDNA120K training set from the single mutant subset that had experimental structures available of the recently published cDNA-display proteolysis Dataset #1 [46] (Figure 2).

**Figure 2:**
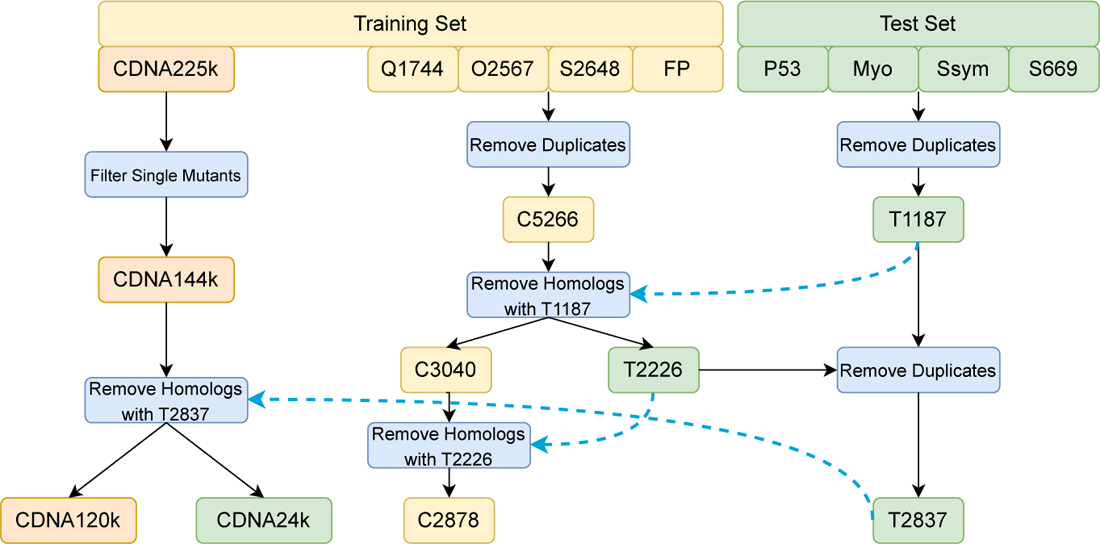
Training and test set generation pipeline. Homologous proteins were removed using MMSeqs2 [45] with a sequence similarity threshold of 30%.

Even with the T2837 expanded test set, we are still unable to assess generalization performance on 14% of the 380 mutation types, since they are not represented in T2837. Additionally, T2837 is heavily biased with mutations to alanine (Figure 3c), further hindering our ability to evaluate the generalization of our model. The community has traditionally relied on the data augmentation technique Thermodynamic Reversibility (TR) [43] to generate datasets with expanded mutation type coverage. However, *∼*3% of mutation types in C2878 + TR and T2837 + TR still lack data (Figure 12 in Appendix A). More importantly, a major drawback of TR augmentation is that all stabilizing mutations it generates are to the wildtype amino acid, as shown in Figure 3b. These mutations give no predictive power with respect to identifying non-wildtype stabilizing mutations, which is the main goal of thermodynamic stability prediction in the context of protein engineering. To improve the predictive power of deep learning frameworks for stabilizing proteins, additional data for stabilizing mutations *not* to the wildtype amino acid is required.

**Figure 3:**
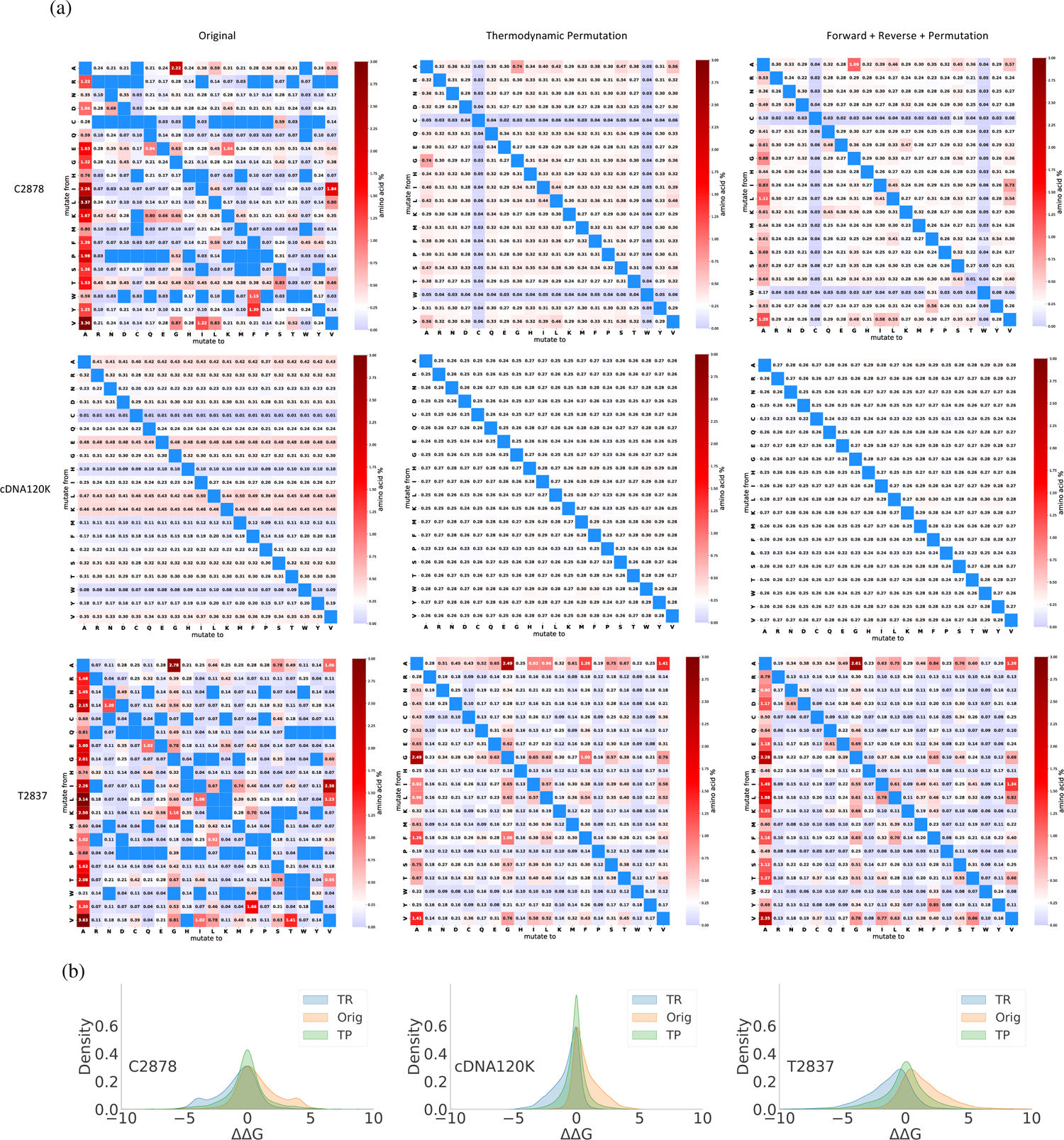
a) Heatmap representation of the mutation type distribution present in C2878, cDNA120K, T2837. For C2878, the original, TP, and original + TP + TR dataset consist of 2878, 18912, and 24668 mutations that sample 86.8%, 100%, and 100% of the mutation types, respectively (first row). For cDNA120K, the original, TP, and original + TP + TR datasets consist of 119690, 2071122, and 2190812 mutations, respectively, and each dataset samples 100% of the mutation types (second row). For T2837, the original, TP, and original + TP + TR datasets consist of 2837, 7720, and 13394 mutations that sample 85.5%, 100%, and 100% of the mutation types, respectively (third row). b) Comparing the ΔΔ*G* distribution for the original training and test sets with their TP and TR augmentations. All three experimental datasets are biased towards destabilizing mutations (orig). TR augmentation provides additional data biased towards stabilizing mutations and TP augmentation provides additional data that is evenly distributed between

To address these issues and improve Stability Oracle’s ability to generalize, we introduce Thermodynamic Permutations (TP), a new data augmentation technique. TP takes advantage of the fact that Gibbs free energy is a state function and enables the generation of thermodynamically accurate point mutations at residues where multiple amino acids have been experimentally characterized. With TP, we can generate an additional 2.1M, 18.9K, and 7.7K point mutations that sample all 380 mutation types for cDNA120K, C2878, and T2837, respectively (Figure 3). Additionally, TP mitigates several sampling biases in all 3 datasets (Figure 3a middle column). First, it provides mutation data for the 13.2% and 14.5% mutation types absent in C2878 and T2837. TP generated data for C2878 and T2837 touches 380 mutation types, providing the first training and test sets with experimental ΔΔ*G* measurements for all mutation types (the cDNA display proteolysis dataset does not directly measure ΔΔ*G* but instead derives ΔΔ*G* values from next-generation sequencing data of proteolytic experiments).

Figure 3a illustrates improvement in sampling bias as a softening of red (oversampled) and blue (undersampled) towards white (balanced sampling). In the C2878 and T2837 datasets, this is most apparent for the “to” alanine bias. In cDNA120K, there is an oversampling bias of mutations “from” alanine, glutamate, leucine, and lysine and an undersampling bias for mutations “from” cysteine, histidine, methionine, and tryptophan. TP completely balances the cDNA120K mutation type distribution with each mutation type making up approximately 0.26% of the dataset (100% / 380), depicted in Figure 3c as uniformly white. Thus, TP augmentation of cDNA120K provides the first large-scale ΔΔ*G* dataset (>1M) that evenly samples all 380 mutation types across 100 protein domains. In contrast to TR, TP does not include stabilizing mutations to wildtype and yields a balanced distribution (stabilizing vs destabilizing) of ΔΔ*G* measurements (Figure 3b).

We fine-tuned Stability Oracle on cDNA120K and C2878 with and without TP augmentation and evaluated its performance on all test sets. As shown in Figure 5a, training on cDNA120K + TP + TR provided the best performance overall on regression and classification metrics across the test sets. While this might have been expected due to the sheer size and mutation-type balance compared to C2878 + TP + TR, it is interesting to note that proteolytic stability of single-domain natural proteins is in fact generally an excellent surrogate for thermodynamic stability (as was pointed out in the original publication [46]). From this data, the impact of TP on model generalization was unclear. To further examine how TP-augmented datasets impact generalization, we evaluated predictions at mutation types in T2837 + TP that were *absent* from C2878 + TR, namely the 12 mutation types with no experimental data (Figure 12 in Appendix A). For these mutation types, TP improves generalization: recall improves from 0.28 to 0.4 and precision improves from 0.467 to 0.667 (Figure 4). We artificially expanded the mutation types that were missing data and observed similar results (Figure 4).

**Figure 4:**
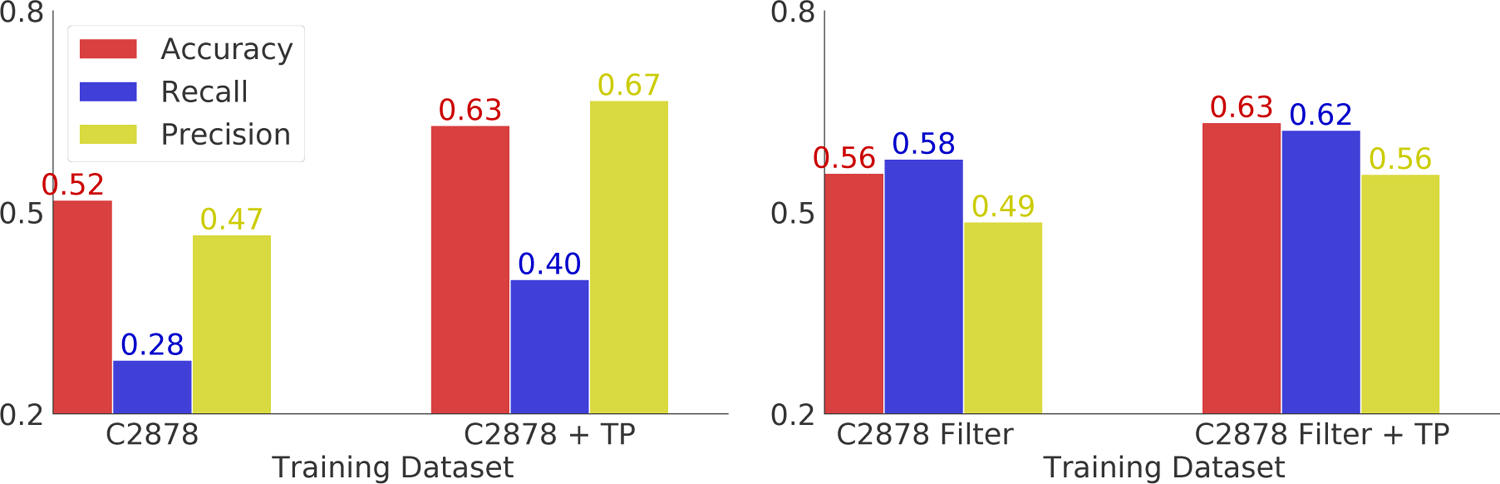
Evaluation of TP on mutation types lacking experimental data in the C2878 training set. We report the accuracy, recall, and precision results on two subsets of T2837 + TP, to demonstrate the effectiveness of permutation. In the left, we test the model on the 12 mutation types lacking experimental data in C2878 + TR. However, this analysis consist of only 54 mutations in T2837 + TP. In the right, we removed mutation types from C2878 that had less than 8 training instances, so 68 mutation types lacked data, and augment this filtered training set with TP. This filtered version of C2878 allowed us to evaluate the performance of TP augmentation on 663 mutations in T2837 + TP.

**Figure 5:**
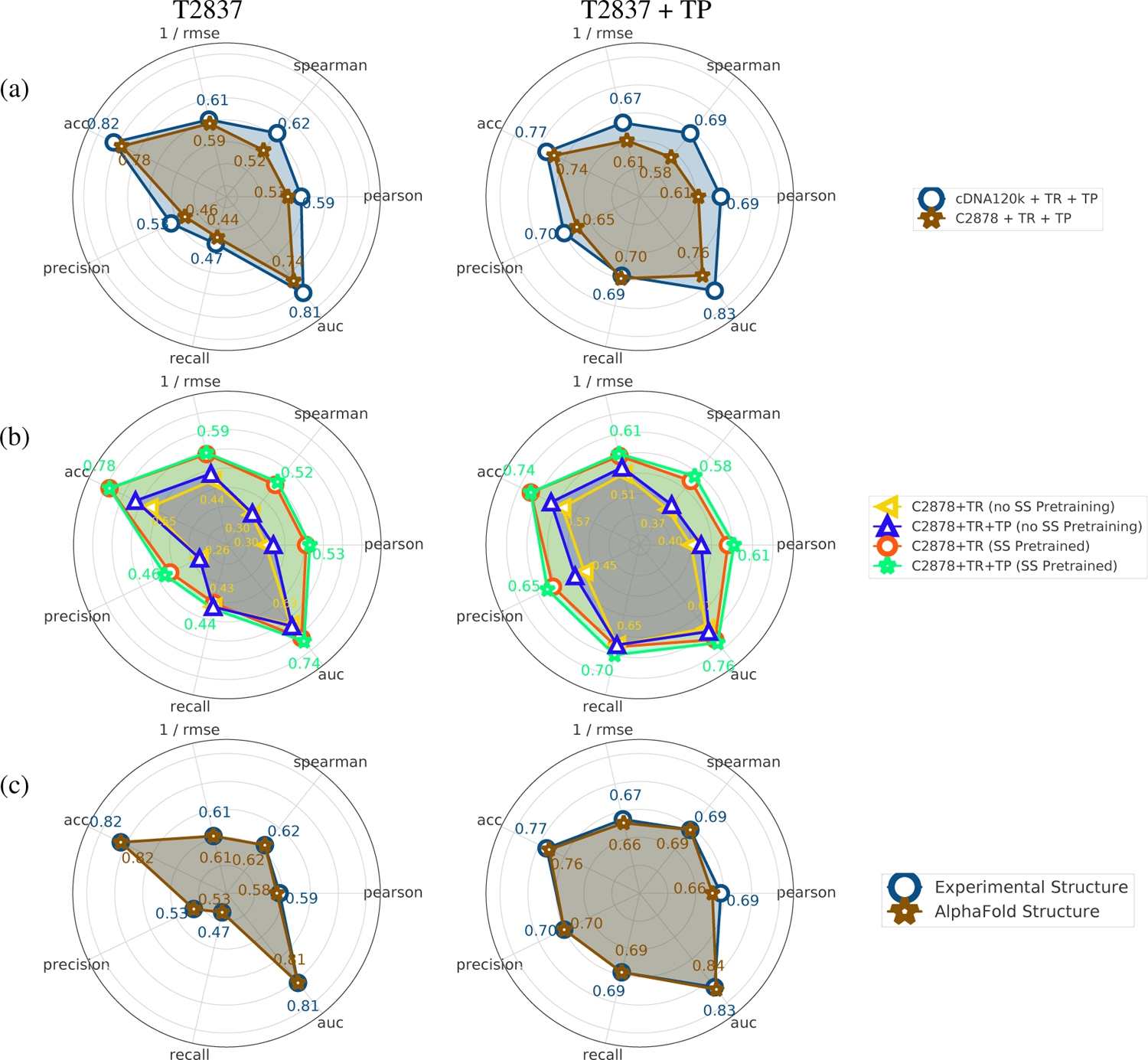
Regression and classification performance of different models on T2837 and T2837TP. (a) Comparison between the C2878 and cDNA120K training sets. (b) Comparison of Stability Oracle’s performance when trained on C2878 with and without a pre-trained backbone and structural amino acid embeddings and with and without TP augmentation. (c) Stability Oracle’s performance when tested on experimental or AlphaFold structures of T2837.

To prevent inflation of Stability Oracle’s binary classification performance, we focus on T2837 + TP and exclude all TR mutations for T2837, since these mutations are heavily biased with stabilizing mutations to the wildtype amino acid (Figure 13 in Appendix A). Here, Stability Oracle demonstrated a recall of 0.69, a precision of 0.70, and an AUROC of 0.83 (Figure 9). Surprisingly, further fine-tuning with C2878 + TP did not improve performance on T2837 or T2837 + TP. Our analysis reveals a 100% homology overlap (>30% sequence similarity) between the proteins in cDNA120K and C2878, providing a rationale for the lack of performance improvement. However, C2878 fine-tuning improves performance on the interface subset of T2837: the Pearson correlation improves from 0.30 to 0.35, and the Spearman correlation improves from 0.29 to 0.35 for the interface microenvironments. This improvement is expected since the cDNA dataset consists of monomeric single-domain structures, lacking interfaces with other proteins, ligands, or nucleotides. However, meaningful improvements is limited due to the scarce amount of protein-protein (127 mutations), protein-ligand (94 mutations), and protein-nucleotide (9 mutations) data in C2878.

Since experimental structures are often unavailable, we examined Stability Oracle’s ability to generalize to structures generated by AlphaFold. We used ColabFold [57] to generate template-free predicted structures for each protein in T2837 and T2837 + TP. ColabFold failed to fold one protein and two structures were removed due to US-align having TM-scores *<* 0.5, which resulted in the removal of 50 mutations from T2837 [58, 59]. We observed no changes in classification metrics and slightly lower performance on regression metrics on T2837 + TP such as Pearson, Spearman, and RMSE (Figure 5c). Overall, these results demonstrate the ability of Stability Oracle to generalize to AlphaFold scaffolds when an experimental structure is unavailable.

To compare against the literature, we report Pearson correlation coefficients (both forward and reverse) on T2837 and all the common test sets. For the common test sets, we compare against several community predictors in Figure 6 and provide additional regression metrics for literature comparisons in Table 3a, Appendix A. It is worth noting that Stability Oracle outperforms other predictors in the literature even with their documented data leakage issues.

**Figure 6:**
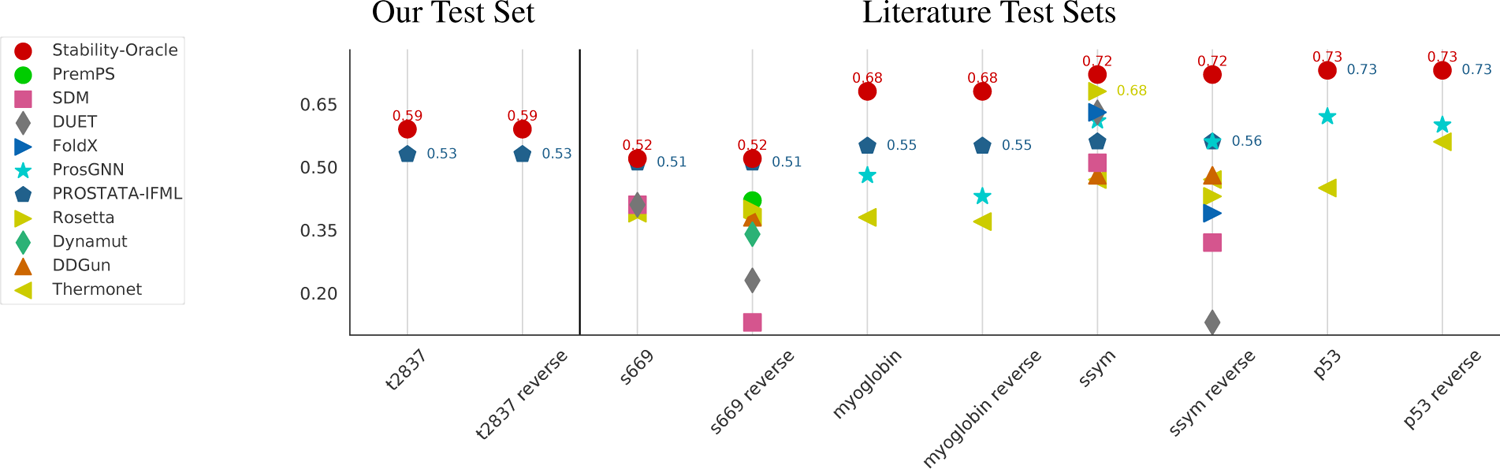
Pearson Correlation Coefficient of Stability Oracle and Prostata-IFML across several test sets. We compare against a handful of computational stability predictors from the community (values obtained from the literature).

### 2.3 Evaluating Stability Oracle’s Ability to Identify Stabilizing Mutations

For computational stability predictors to accelerate protein engineering, it is critical that their predictions correctly identify stabilizing mutations. However, it is well documented that state-of-the-art stability predictors can correctly predict stabilizing mutations with only a 20% success rate and most stabilizing predictions are actually neutral or destabilizing [31, 13]. While molecular dynamic-based methods, such as Free Energy Perturbation (FEP), have demonstrated a 50% success rate at identifying stabilizing mutations, their computational demand prevents them from scaling to entire protein applications like computational deep mutational scans (DMS) [31, 60]. Thus, there is a strong need for a method that can match the performance of FEP while being computationally inexpensive.

To evaluate Stability Oracle’s ability to identify stabilizing mutations, we filtered its predictions on T2837 and T2837 + TP at different ΔΔ*G* thresholds and assessed the distribution of experimental stabilizing (ΔΔ*G* < −0.5 kcal/mol), neutral (|ΔΔ*G*| *≤* 0.5 kcal/mol), and destabilizing (ΔΔ*G* > 0.5 kcal/mol) mutations. The 0.5 kcal/mol cutoff was chosen based on the average experimental error [61]. With the ΔΔ*G* < −0.5 kcal/mol prediction threshold, 1770 mutations were filtered with an experimental distribution of 74.0% stabilizing, 17.8% neutral, and 8.2% destabilizing and 48.1% of all stabilizing mutations were correctly identified. A systematic analysis of prediction thresholds are provided in the Appendix (Figure 10 and Table 4a). The success rate of stabilizing mutations surpasses what is typically observed with FEP methods (*∼*50%) [31, 60] with orders of magnitude less computational cost (Table 1, Appendix A). We further examine Stability Oracle’s ability to identify stabilizing mutations by amino acid (Figure 7). Here, we observe that Stability Oracle is able to correctly predict stabilizing mutations across most amino acids, whether mutating “from” or “to”. However, several amino acids lack sufficient “from” or “to” stabilizing predictions to draw meaningful conclusions. This data scarcity is even more apparent when looking at the 380 “from”-“to” pairs (see Figure 14, Appendix A), highlighting how data scarcity still hinders proper model evaluation.

**Figure 7:**
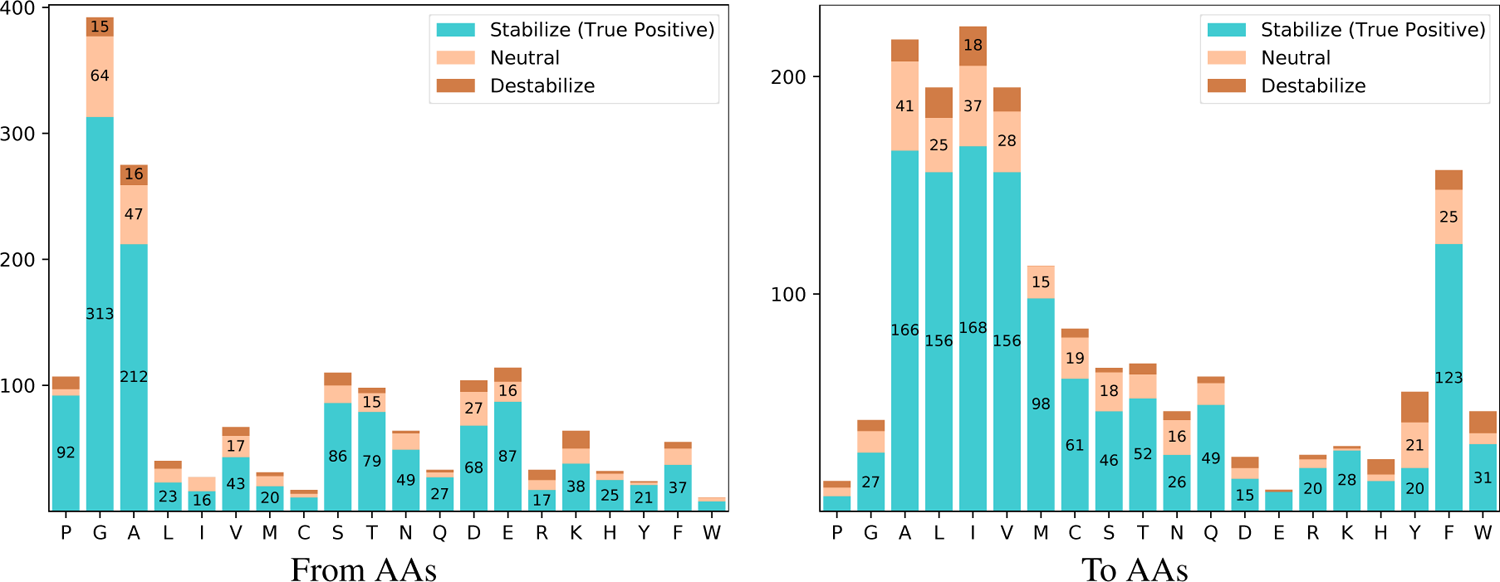
Experimental distribution of Stability Oracle’s stabilizing predictions (ΔΔ*G* < −0.5 kcal/mol) on T2837 + TP test set for different “from” and “to” amino acid types. Here, stabilizing, neutral, and destabilizing mutations are defined by ΔΔ*G* < −0.5 kcal/mol, |ΔΔ*G*| *≤* 0.5 kcal/mol, and ΔΔ*G* > 0.5 kcal/mol, respectively.

**Figure 8:**
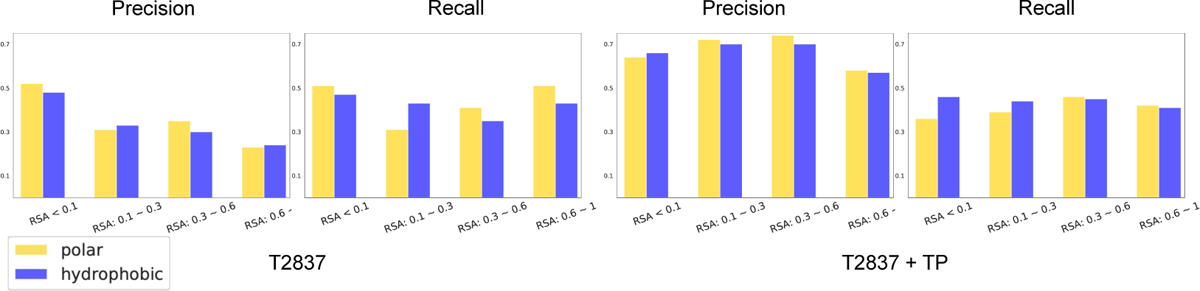
Comparing Stability Oracle’s precision and recall between hydrophobic and hydrophilic amino acids as a function of relative solvent accessibility (RSA). The results demonstrate that there are no biases between polar and hydrophobic residues throughout a protein structure on both T2837 and T2837 + TP.

**Figure 9:**
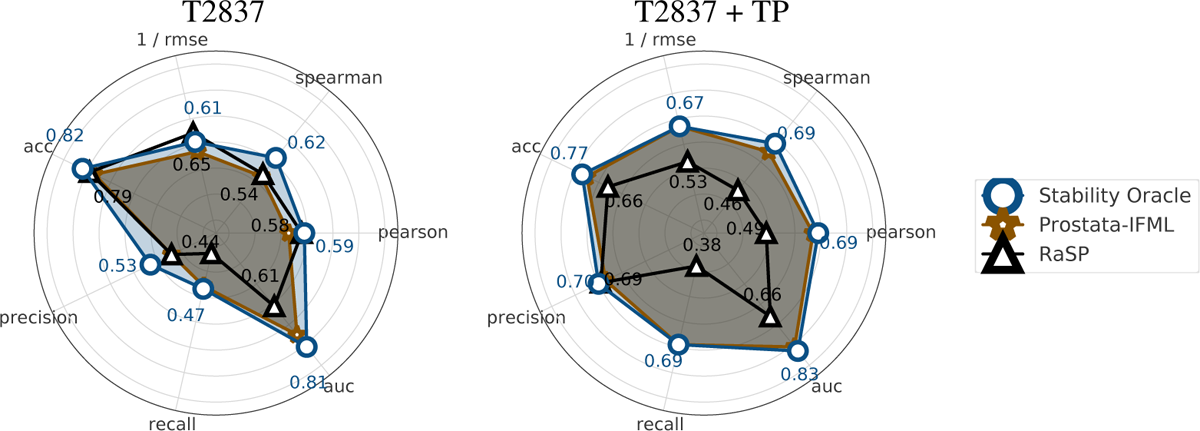
The Comparison of Stability Oracle, Prostata-IFML and RaSP regression and classification performance of different models on T2837 and T2837 + TP. We refer the readers to Appendix A for detailed results.

**Figure 10:**
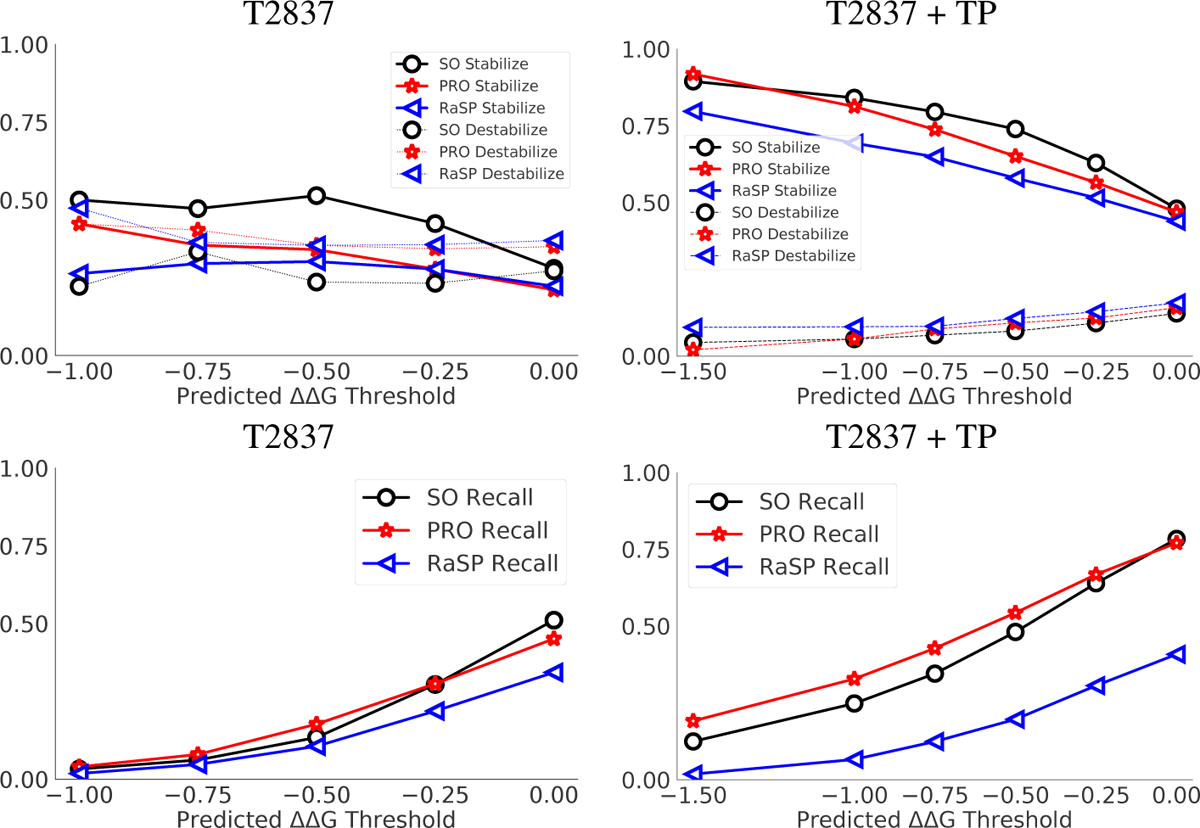
We compare SO (Stability Oracle) with PRO (Prostata-IFML) and RaSP. In the first column, we compare on T2837 and observe that Stability Oracle had the highest stabilization and lowest destabilization fraction with similar recall. In the second column, we compare on T2837 + TP and observe that Stability Oracle still has the highest stabilization and lowest destabilization fraction but Prostata-IFML has better recall.

It has been pointed out by the community that the subset of experimentally validated stabilizing surface mutations are heavily skewed towards mutations to hydrophobic amino acids. An analysis by Broom et al. of the ProTherm database [62] indicates that surface stabilizing mutation bias typically increases side chain hydrophobicity (ΔΔ*G*_solvation_) with a median change of 0.8 kcal/mol. This hydrophobicity bias is equivalent to an alanine-to-valine mutation on the protein surface [31]. We examined if this underlying hydrophobic bias persisted within our training pipeline by computing the precision and recall of polar and hydrophobic mutations as a function of relative solvent accessibility (RSA) of the wildtype residue. Our precision and recall results across different RSA bins of T2837 and T2837+ TP indicate that there is not a bias between stabilizing predictions to hydrophobic and polar amino acids across a protein structure (see Figure 8). This demonstrates that the cDNA120K + TP training set mitigates a well known bias in computational stability prediction training sets.

### 2.4 Comparing Sequence and Structure Fine-Tuned Stability Predictors

Over the last three years, self-supervised protein Large Language Models (pLLMs or “sequence models”) have had a tremendous impact on the protein community [63, 64, 65, 66, 67, 68, 69, 70, 25].

Understanding sequence versus structure-based prediction models continues to be an active area of research in protein engineering and design [71, 19]. We evaluated Stability Oracle against two computational stability deep learning frameworks: Prostata and RaSP. Prostata [26] is a sequence-based framework that fine-tunes ESM2 embeddings (a SOTA pLLM) [25] on common training and test sets. To address known data leakage and conduct a fair comparison, we fine-tuned Prostata models on the same training and test sets as Stability Oracle. We call our version of this sequence model Prostata-IFML. We also compare against the RaSP framework [72]: a structure-based CNN model that follows a similar training pipeline. Briefly, RaSP is pre-trained with 2315 structures clustered at a 30% sequence similarity and then fine-tuned on 35 DMS datasets computationally generated by the Rosetta Cartesian-ΔΔ*G* program. Like Stability Oracle, RaSP uses a masked microenvironment paradigm to avoid the need to generate mutant structures. In contrast to Stability Oracle, RaSP represents the masked microenvironments as an 18Å voxelized cube and the mutations via one-hot encodings of the “from” and “to” amino acids. In our analysis, we modified the RaSP Colab notebook to generate DMS predictions on every protein in T2837 and every mutation in T2837 + TP.

Stability Oracle outperforms or matches Prostata-IFML on every metric (Figure 9), even though Stability Oracle has 548 times fewer parameters (*∼*658.M vs *∼*1.2M) and was pre-trained with 2,000 times fewer proteins (*∼*46M vs *∼*23K) at the same sequence similarity (50%). As for RaSP, Stability Oracle significantly outperforms it on nearly every classification and regression metric on both T2837 and T2837 + TP. The performance is only comparable for Pearson on T2837 and precision on T2837 + TP.

In terms of identifying stabilizing mutations, Stability Oracle also achieves the best performance (Figure 10). At each prediction threshold, Stability Oracle had the highest proportion of correctly identified stabilizing mutations and the lowest proportion of destabilizing mutations (we exclude −1.5 kcal/mol threshold for T2837 due to data scarcity). It is typical for precision to be inversely proportional to recall, and we observe this tradeoff with regard to Stability Oracle and Prostata-IFML on T2837 + TP, where Prostata-IFML has better recall. We suspect this difference is due to their loss: Stability Oracle uses a huber loss and Prostata uses mean square error. Nonetheless, both Stability Oracle and Prostata-IFML have superior performance on both on correctly identifying stabilizing mutations (precision) and identifying more stabilizing mutations (recall) compared to RaSP. A detailed comparison is provided in Appendix A.8.

Finally, we compare all three framework’s ability to predict self-mutations. Here, the “from” and “to” amino acid are the same and the ΔΔ*G* should equal 0 kcal/mol. This experiment is similar to the forward vs reverse experiments and is meant to assess the robustness of the predictors. For wildtype self-mutations on T2837 Stability Oracle, Prostata-IFML, and RaSP achieve RMSE of 0.0033, 0.0018, and 0.8370 kcal/mol, respectively. This demonstrates that the training pipeline for Stability Oracle and Prostata-IFML implicitly captures self-mutations. However, RaSP is unable to generalize to self-mutations and this drop in performance is also observed for TR augmentation of T2837 (Appendix A.9).

## 3 Discussion

Reliable prediction of stabilizing mutations is critical for the acceleration of protein-based biotechnologies. However, to date, all computational stability predictors prioritize improvements in the Pearson and RMSE metrics. Neither of these metrics, however, is appropriate for evaluating improvements in identifying stabilizing mutations; several studies explain this in great detail [31, 14, 13]. We report these regression metrics but use classification metrics, such as AUROC, MCC, recall, and precision, to guide our hyperparameter tuning. By using a 30% sequence similarity threshold for train-test splits, we expect the performance of Stability Oracle and Prostata-IFML to have superior generalization compared to models trained using traditional train-test splits, which suffer from data leakage. Stability Oracle and Prostata-IFML seem to correctly identify stabilizing mutations that potentially outperform FEP-based methods while being several orders of magnitude faster. However, experiments that directly compare FEP-based methods are needed for confirmation.

The literature has recently adopted Thermodynamic Reversibility (TR) to address the imbalance between stabilizing and destabilizing mutations in the training and test sets. However, this biases the training towards stabilizing mutations of wildtype amino acids. For computational models that leverage evolutionary features and protein structures as input, mutations to wildtype amino acids leak information and are of limited use in a protein engineering context where mutating away from wildtype is the primary goal. Our data augmentation technique, Thermodynamic Permutations (TP), generates mutations with a balanced ΔΔ*G* distribution and does not generate mutations to wildtype amino acids (see Figure 3b and Figure 13 in Appendix A). This mitigates the above imbalance and expands the number of stabilizing mutations to non-wildtype amino acids in both the training and test sets. In addition, TP reduces bias to wildtype inherent in the self-supervised pre-training step. We speculate that during end-to-end fine-tuning, TP forces the feature extractor to find additional chemical signatures in the microenvironment, not just those most relevant for wildtype identification. Further, TP also generates ΔΔ*G* measurements for mutation types in microenvironments that would rarely be experimentally characterized. This expands the microenvironment/mutation type combinations available for training and testing and is highly likely to improve generalization across structural motifs found within a protein scaffold. However, we lack the necessary test data needed to examine this hypothesis. We expect TP to be of great use for the development of frameworks for higher-order mutations, where data scarcity is an even bigger issue.

Of note is the performance of Stability Oracle’s much smaller model (in terms of parameters) relative to Prostata-IFML. Stability Oracle’s ability to outperform or match Prostata-IFML is evidence that structures contain information beyond the amino acid sequence itself. Self-supervised deep learning models, such as MutComputeX, typically struggle with predicting mutations in the core, with a tendency to re-predict the wild-type residue in the closely packed environment. Stability Oracle is explicitly trained to learn stabilizing substitutions from the microenvironment, providing a structure-based method for predicting core mutations.

In terms of training set size, note that the *∼*25,000 ΔΔ*G* measurements (C2878 + TP + TR) led to performance comparable to models trained on *∼*2.2 million proteolytic stability measurements (cDNA120K + TP + TR). This indicates that both the quality and quantity of data is of great importance for training. Additional experiments directed towards producing large datasets with detailed thermodynamic information, especially for residues at functional interfaces, may further improve generalization capabilities.

Accurate identification of stabilizing mutations will continue to impact a wide variety of areas, from predicting protein therapeutics and vaccines with greater stabilities and shelf lives, to predicting enzymes that can work at higher temperatures for bioprocessing and environmental bioremediation. While previous algorithms, such as MutCompute, have proven adept at improving the stabilities of proteins [47, 48, 50], Stability Oracle will likely further improve the hit rate of functional, stable substitutions due to its improved accuracy across a variety of metrics. More importantly, Stability Oracle is attuned to thermodynamic effects, rather than just sterics. This allows for predictions across a wider class of substitutions, including those at protein:protein interfaces, such as antibody:antigen interactions, as well as those at protein:ligand and protein:nucleotide interfaces.

## Acknowledgments

This work was sponsored by grants from the Defense Threat Reduction Agency (HDTRA1201001) and the Welch Foundation (F-1654) and the NSF AI Institute for Foundations of Machine Learning (IFML). We would like to thank AMD for the donation of critical hardware and support resources from its HPC Fund.

## A Supplementary Data

### A.1 Running time on different proteins

**Table 1:** We present the time to generate the masked microenvironment graph per residue (data processing time) and predict the ΔΔ*G* for the 20 amino acids per residue (inference time). These performance benchmarks were on one 2.00 GHz Intel(R) Xeon(R) Gold 6338 CPU core and model inference time on one A100 GPU.

| PDB ID | # Residues | Per-Residue Graph Generation Time (s) | Per-Residue Inference Time (s) |
| --- | --- | --- | --- |
| 4DCH | 217 | $0.0193 \pm 0.0001$ | $0.0783 \pm 0.0042$ |
| 4AJY | 355 | $0.0108 \pm 0.0001$ | $0.0609 \pm 0.0012$ |
| 2R7E | 1337 | $0.0273 \pm 0.0001$ | $0.0457 \pm 0.0001$ |
| 6VSB | 2905 | $0.0514 \pm 0.0001$ | $0.0422 \pm 0.0004$ |

### A.2 Evaluating our structural amino acid embeddings v.s. one-hot encodings

**Table 2:**
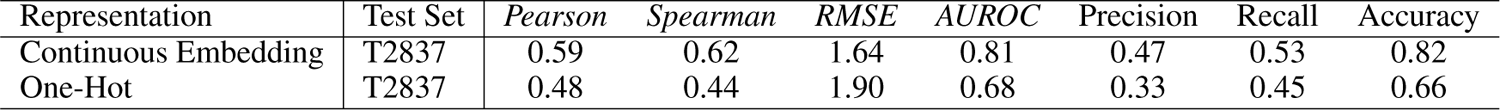
We present the comparison of two different protein sequence representations, Continuous Embedding, and One-Hot, on the T2837 test set. The table shows the average values of the Pearson correlation coefficient, Spearman correlation coefficient, Root Mean Squared Error (RMSE), Area Under the Receiver Operating Characteristic Curve (AUROC), Precision, Recall, and Accuracy. Results are the average of three independently trained models on the cDNA120K dataset.

| Representation | Test Set | *Pearson* | *Spearman* | *RMSE* | *AUROC* | Precision | Recall | Accuracy |
| --- | --- | --- | --- | --- | --- | --- | --- | --- |
| Continuous Embedding | T2837 | 0.59 | 0.62 | 1.64 | 0.81 | 0.47 | 0.53 | 0.82 |
| One-Hot | T2837 | 0.48 | 0.44 | 1.90 | 0.68 | 0.33 | 0.45 | 0.66 |

### A.3 Diagram describing the TR and TP augmentations

**Figure 11:**
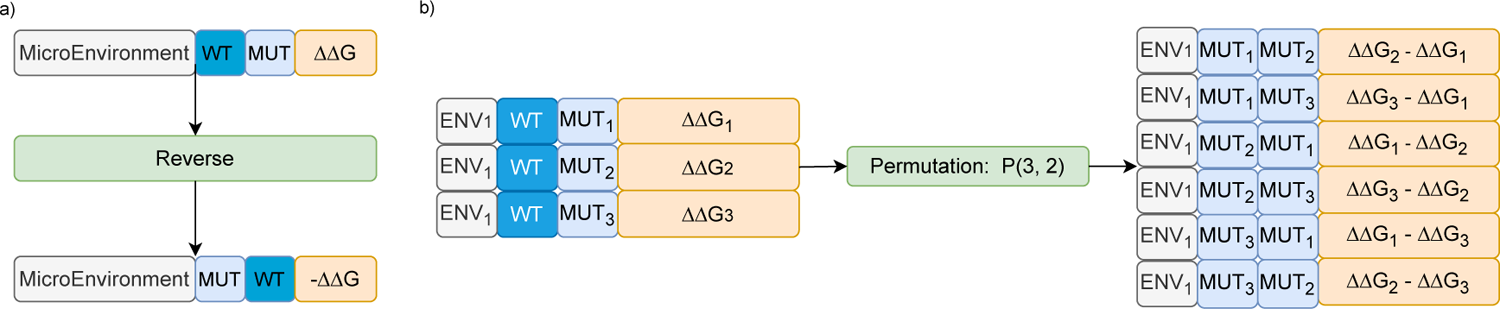
Thermodynamic data augmentation techniques applied to our datasets. a) Diagram describing how Thermodynamic Reversibility works (TR) b) Diagram describing how Thermodynamic Permutations works (TP). Note: TP augmented data does not include the wildtype amino acid.

### A.4 Distribution of experimental datasets by mutation type with Thermodynamic Reversibility

**Figure 12:**
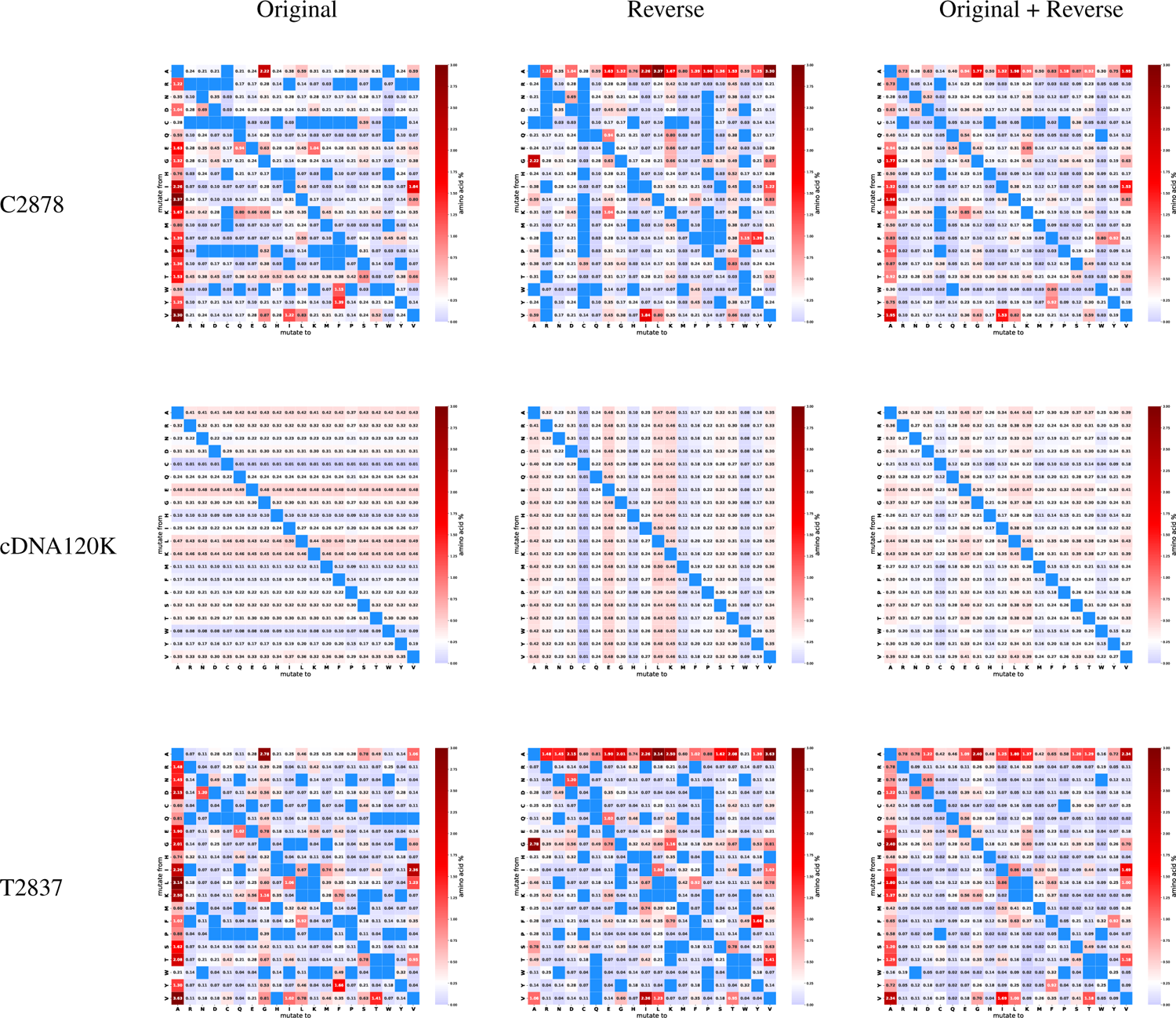
Heatmap of C2878, cDNA120K, and T2837. First column shows the original data, middle column shows the TR augmented data, and right column shows the original + TR datasets.

### A.5 Distribution of experimental datasets by mutation type with Thermodynamic Permutations

**Figure 13:**
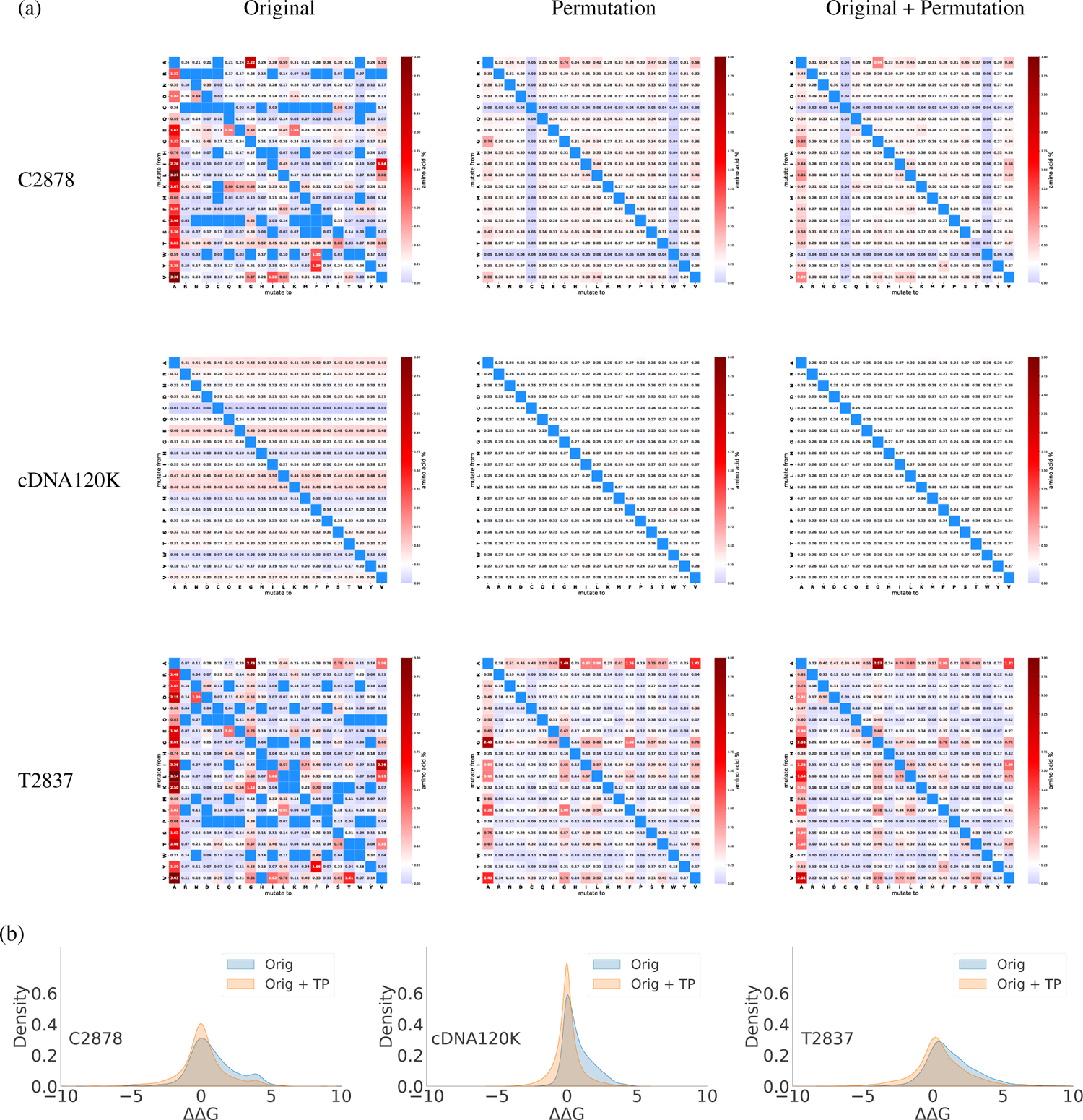
(a) Heatmap of C2878, cDNA120K, and T2837. The first column shows the original data, the middle column shows the TP augmented data, and the right column shows the original + TP datasets. (b) Density plots that superimpose the original datasets and the TP augmented dataset. This plot highlights how TP augmentation adds stabilizing mutations that do not include the wildtype amino acids.

### A.6 Stabilization ratio for the 380 mutation types

**Figure 14:**
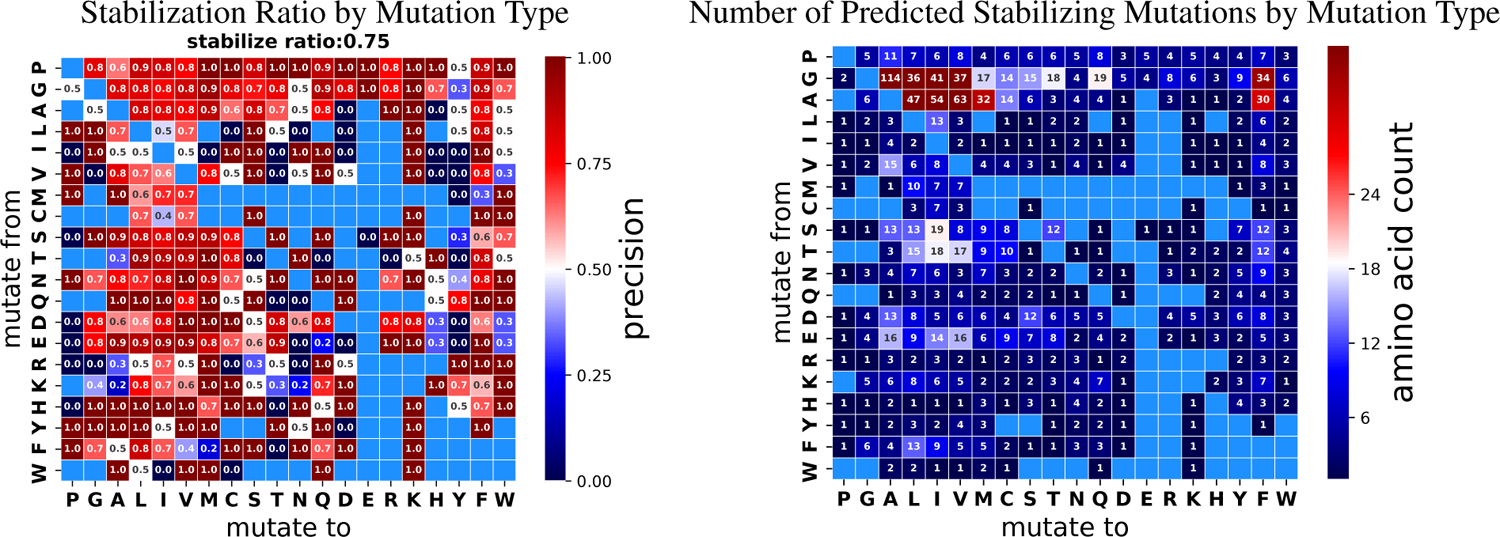
Distribution of Stability Oracle’s stabilizing predictions with a stability threshold of ΔΔ*G < −*0.5 kcal/mol. Left shows the ratio of predictions that are experimentally stabilizing for each mutation type. Right shows the number of mutations predicted to be stabilizing for each mutation type, highlighting the lack of data for evaluation.

### A.7 Performance of Stability Oracle and Prostata-IFML on Common Test Sets

**Table 3:**
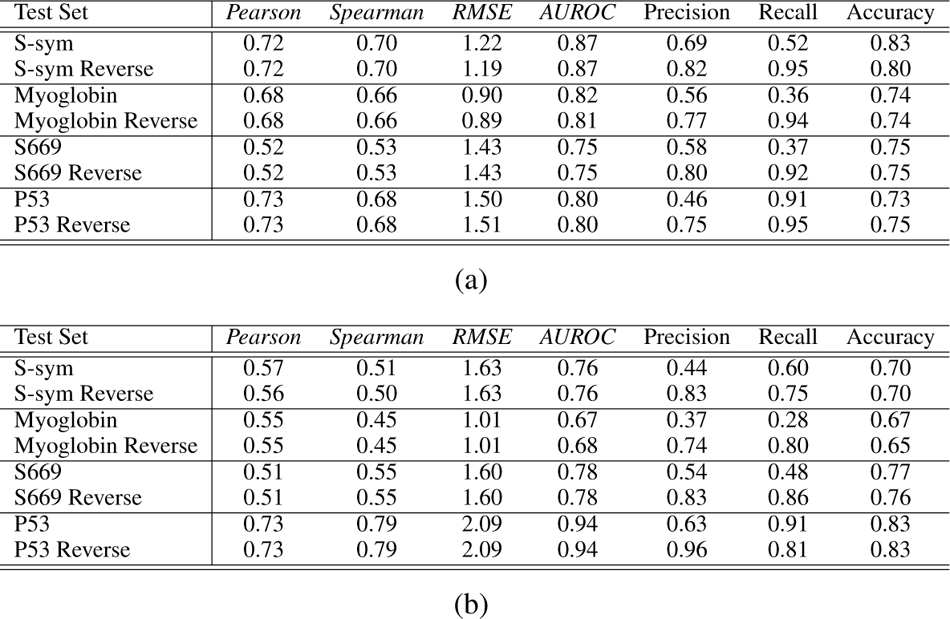
In (a), we present the performance of the Stability Oracle on various test sets. We report the Pearson and Spearman correlation coefficients, root mean square error (RMSE), the area under the receiver operating characteristic (AUROC) curve, Precision, Recall, and Accuracy. In (b), we present the performance of PROSTATA-IFML on various test sets. We report the Pearson and Spearman correlation coefficients, root mean square error (RMSE), the area under the receiver operating characteristic (AUROC) curve, Precision, Recall, and Accuracy.

| Test Set | Pearson | Spearman | RMSE | AUROC | Precision | Recall | Accuracy |
| --- | --- | --- | --- | --- | --- | --- | --- |
| S-sym | 0.72 | 0.70 | 1.22 | 0.87 | 0.69 | 0.52 | 0.83 |
| S-sym Reverse | 0.72 | 0.70 | 1.19 | 0.87 | 0.82 | 0.95 | 0.80 |
| Myoglobin | 0.68 | 0.66 | 0.90 | 0.82 | 0.56 | 0.36 | 0.74 |
| Myoglobin Reverse | 0.68 | 0.66 | 0.89 | 0.81 | 0.77 | 0.94 | 0.74 |
| S669 | 0.52 | 0.53 | 1.43 | 0.75 | 0.58 | 0.37 | 0.75 |
| S669 Reverse | 0.52 | 0.53 | 1.43 | 0.75 | 0.80 | 0.92 | 0.75 |
| P53 | 0.73 | 0.68 | 1.50 | 0.80 | 0.46 | 0.91 | 0.73 |
| P53 Reverse | 0.73 | 0.68 | 1.51 | 0.80 | 0.75 | 0.95 | 0.75 |

(a)
| Test Set | Pearson | Spearman | RMSE | AUROC | Precision | Recall | Accuracy |
| --- | --- | --- | --- | --- | --- | --- | --- |
| S-sym | 0.57 | 0.51 | 1.63 | 0.76 | 0.44 | 0.60 | 0.70 |
| S-sym Reverse | 0.56 | 0.50 | 1.63 | 0.76 | 0.83 | 0.75 | 0.70 |
| Myoglobin | 0.55 | 0.45 | 1.01 | 0.67 | 0.37 | 0.28 | 0.67 |
| Myoglobin Reverse | 0.55 | 0.45 | 1.01 | 0.68 | 0.74 | 0.80 | 0.65 |
| S669 | 0.51 | 0.55 | 1.60 | 0.78 | 0.54 | 0.48 | 0.77 |
| S669 Reverse | 0.51 | 0.55 | 1.60 | 0.78 | 0.83 | 0.86 | 0.76 |
| P53 | 0.73 | 0.79 | 2.09 | 0.94 | 0.63 | 0.91 | 0.83 |
| P53 Reverse | 0.73 | 0.79 | 2.09 | 0.94 | 0.96 | 0.81 | 0.83 |

### A.8 Performance on identifying stabilizing mutations

**Table 4:**
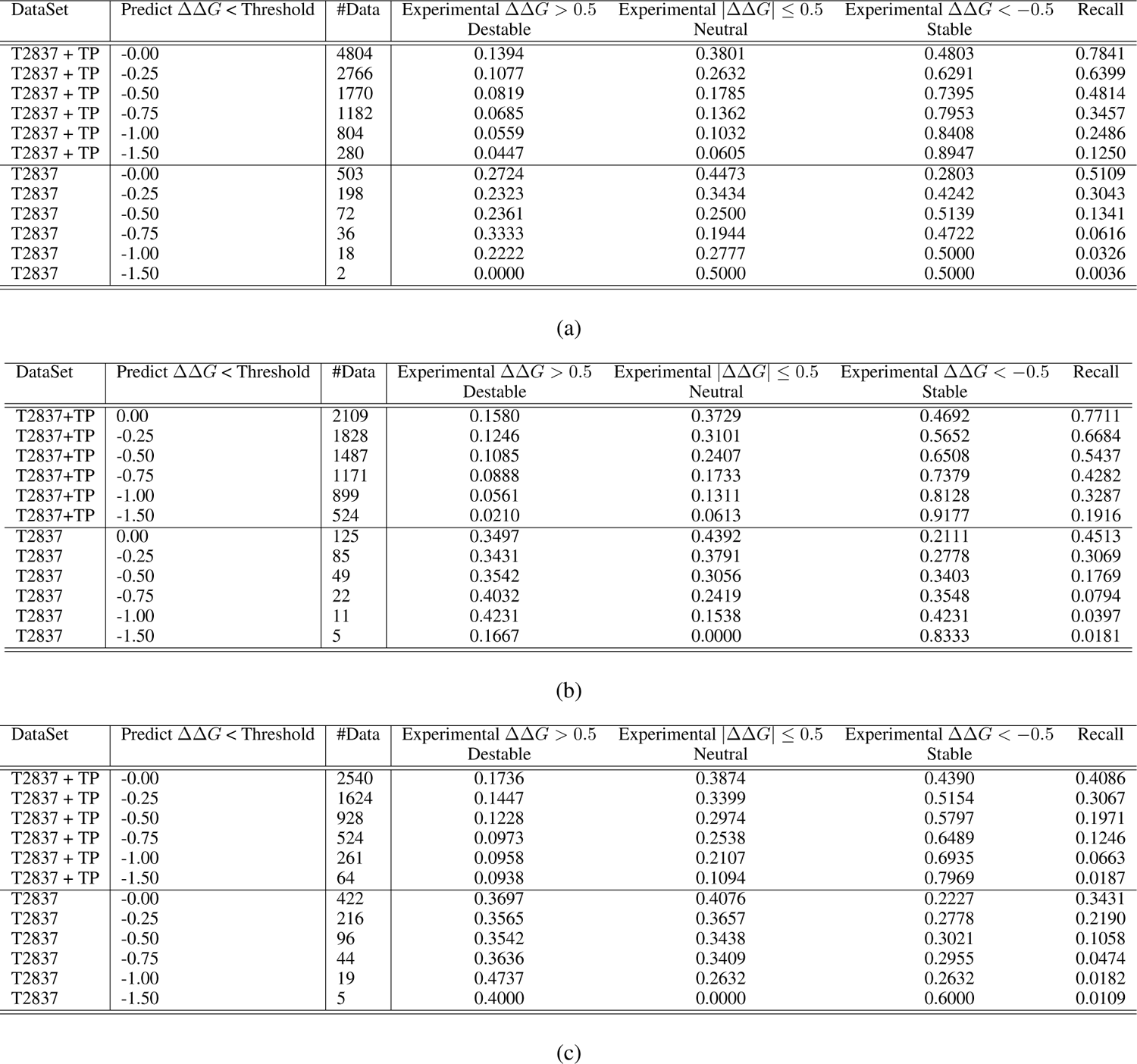
(a) The experimental distribution of Stability Oracle on T2837 and T2837 + TP with different prediction ΔΔ*G* thresholds. (b) The experimental distribution of Prostata-IFML on T2837 and T2837 + TP with different prediction ΔΔ*G* thresholds. (c) The experimental distribution of RaSP on T2837 and T2837 + TP with different prediction ΔΔ*G* thresholds.

### A.9 Regression and classification performance on T2837 and its augmentations

**Table 5:** (a) Stability Oracle’s regression and classification metrics on T2837 and its augmented datasets. TR: Thermodynamic Reversibility augmentation, TP: Thermodynamic Permutation augmentation. Note: mutations from proteins that failed the data engineering pipeline are excluded. (b) Prostata-IFML’s regression and classification metrics on T2837 and its augmented datasets. TR: Thermodynamic Reversibility augmentation, TP: Thermodynamic Permutation augmentation. (c) RaSP’s regression and classification metrics on T2837 and its augmented datasets. TR: Thermodynamic Reversibility augmentation, TP: Thermodynamic Permutation augmentation. Note: we cannot calculate MCC and AUROC for the experimental stable dataset since there is only a single label. Note: mutations from proteins that failed the data engineering pipeline are excluded.

| Test Set | #Mutations | *Pearson* | *Spearman* | *RMSE* | *MCC* | *AUROC* | Precision | Recall | Accuracy |
| --- | --- | --- | --- | --- | --- | --- | --- | --- | --- |
| T2837 Orig | 2830 | 0.5874 | 0.6216 | 1.6549 | 0.3897 | 0.8088 | 0.5381 | 0.4677 | 0.8159 |
| T2837 TR | 2830 | 0.5904 | 0.6226 | 1.6464 | 0.3905 | 0.8138 | 0.8619 | 0.9186 | 0.8159 |
| T2837 Orig + TR | 5660 | 0.7426 | 0.7808 | 1.6527 | 0.6345 | 0.8951 | 0.8074 | 0.8290 | 0.8171 |
| T2837 TP | 7698 | 0.6556 | 0.6500 | 1.4359 | 0.4742 | 0.8107 | 0.7287 | 0.7410 | 0.7371 |
| T2837 Orig + TP | 10528 | 0.6829 | 0.6839 | 1.5300 | 0.5042 | 0.8356 | 0.6903 | 0.6787 | 0.7704 |
| T2837 Orig + TP + TR | 13358 | 0.7034 | 0.7156 | 1.5313 | 0.5399 | 0.8486 | 0.7597 | 0.7830 | 0.7698 |

(a)
| Test Set | Mutations | *Pearson* | *Spearman* | *RMSE* | *MCC* | *AUROC* | Precision | Recall | Accuracy |
| --- | --- | --- | --- | --- | --- | --- | --- | --- | --- |
| T2837 Orig | 2837 | 0.5285 | 0.5284 | 1.7467 | 0.3120 | 0.7469 | 0.4392 | 0.4643 | 0.7726 |
| T2837 TR | 2837 | 0.5289 | 0.5300 | 1.7464 | 0.3236 | 0.7502 | 0.8607 | 0.8634 | 0.7772 |
| T2837 Orig + TR | 5674 | 0.6948 | 0.7026 | 1.7465 | 0.5577 | 0.8513 | 0.7733 | 0.7840 | 0.7749 |
| T2837 TP | 7720 | 0.6788 | 0.6436 | 1.3750 | 0.4473 | 0.8018 | 0.7161 | 0.7315 | 0.7185 |
| T2837 Orig + TP | 10557 | 0.6708 | 0.6484 | 1.4841 | 0.4625 | 0.8093 | 0.6796 | 0.6973 | 0.7331 |
| T2837 Orig + TP + TR | 13394 | 0.6827 | 0.6704 | 1.5433 | 0.4941 | 0.8229 | 0.7403 | 0.7538 | 0.7424 |

(b)
| Test Set | #Mutations | *Pearson* | *Spearman* | *RMSE* | *MCC* | *AUROC* | Precision | Recall | Accuracy |
| --- | --- | --- | --- | --- | --- | --- | --- | --- | --- |
| T2837 Orig | 2830 | 0.5675 | 0.5415 | 1.5557 | 0.2549 | 0.6143 | 0.4384 | 0.3327 | 0.7852 |
| T2837 TR | 2830 | 0.2326 | 0.2733 | 2.3971 | 0.1613 | 0.5999 | 0.8645 | 0.5129 | 0.5484 |
| T2837 Orig + TR | 5660 | 0.551 | 0.5372 | 2.0207 | 0.3573 | 0.6653 | 0.7622 | 0.4772 | 0.6668 |
| T2837 TP | 7714 | 0.4663 | 0.4145 | 1.9767 | 0.2421 | 0.6081 | 0.6917 | 0.3837 | 0.6103 |
| T2837 Orig + TP | 10544 | 0.4913 | 0.4588 | 1.8730 | 0.2684 | 0.6165 | 0.6496 | 0.3772 | 0.6572 |
| T2837 Orig + TP + TR | 13374 | 0.4934 | 0.4660 | 1.9954 | 0.2916 | 0.6323 | 0.7237 | 0.4233 | 0.6342 |

### A.10 The distribution for Stability Oracle’s predicted ΔΔ*G* v.s. experimental ΔΔ*G*

**Figure 15:**
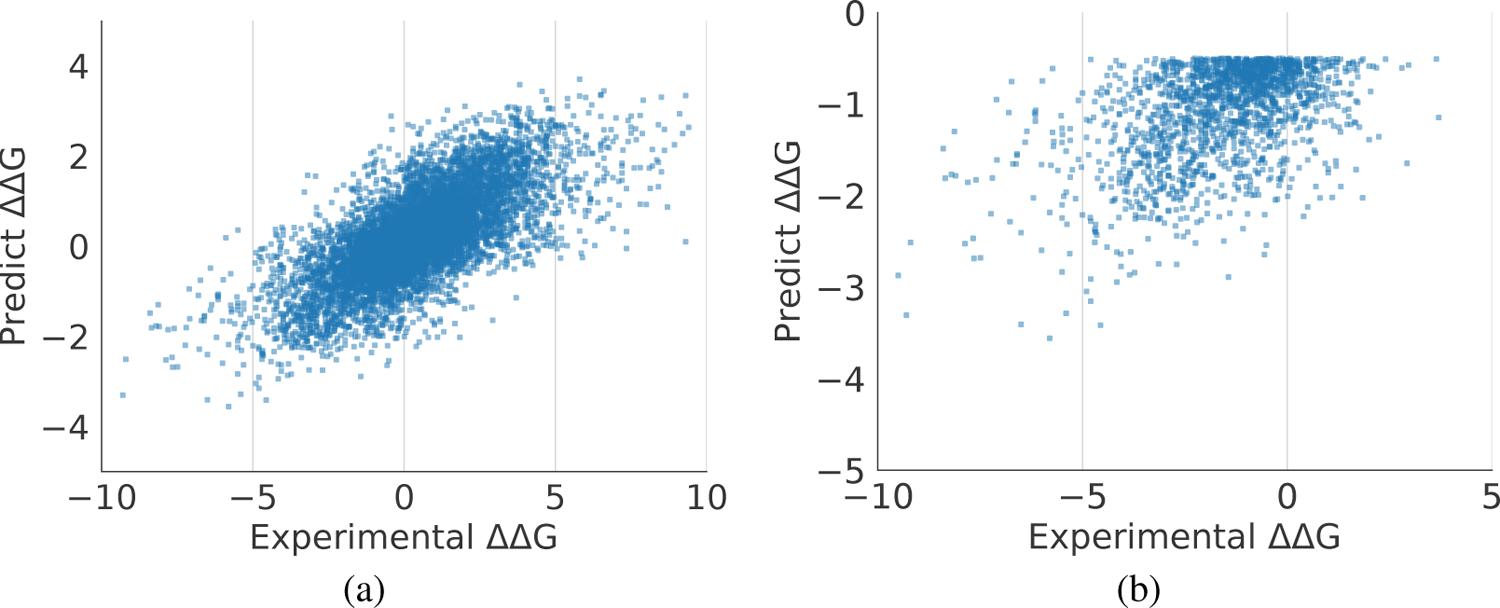
Stability Oracle’s ΔΔ*G* predictions vs experimental ΔΔ*G* measurements.

## B Experiment Details

### B.1 Model Architecture

We build a transformer-based neural network for both mask amino acid self-supervised learning and ΔΔ*G* supervised learning. We describe the model architectures for the self-supervised backbone model and for the ΔΔ*G* regression head.

### Graph Transformer for Self-Supervised Pre-training

We first introduce our graph transformer model for self-supervised tasks, outlining its key components and specifications. We describe the general pictures of our model, and then we list the model inputs and outputs. Finally, we elaborate some key inductive bias designs in our model. Generally speaking, in our tasks, the input for the model is a local environment for the target amino acid. Given one amino acid, we set the Carbon-*α* as the center and grab all the atoms within radius *n*. In the context of self-supervised training, we apply masking to each atom within the target amino acid and predict the corresponding amino acid type. This approach leverages graph-based representations and self-supervised learning to capture important structural features and enhance our understanding of protein sequences.

Denote *N* as the number of tokens, the coordinates *∈* R*^N×^*^3^, atom types *∈* R*^N×^*^1^ and physical properties *p ∈* R*^N×P^* are given to the neural network, we apply an embedding layer to convert categorical atom types into continuous representations *e ∈* R*^N×E^*, and concatenate embeddings together with physical properties *h* = Concat(*e, p*). The coordinates *∈* R*^N×^*^3^ are used to calculate atom-wise euclidean distance *D ∈* R*^N×N^*, which serves as the attention bias in the attention layers. The concatenated features *h* then pass through several attention blocks. In each attention block, we have two attention layers and one MLP layers. The MLP layers is the same as the standard attention block, while in each attention layer, after calculating the *KQ^T^*, we add additional attention mask and attention bias terms.

The attention bias is calculated based on the distance matrix *D ∈* R*^N×N^*. Given *D*, we create multiple feature vectors and output the attention bias for each head in one attention layer. Apply *K* = 16 radial basis functions (RBF) with different bandwidth, we get *k ∈* R*^N×N×K^*. Categorized the distance into *C* = 4 class, we get *c ∈* R*^N×N×C^*. Put a linear transformation after concatenating *c* and *k*, we get bias *∈* R*^N×N×^*^Head^ where Head denotes number of heads in one attention layer.

Finally, we collect the hidden representations *h* for all the atoms, and select those whose corresponding atom is close to the masked amino acid. Specifically, we set the carbon-*α* for the masked amino acid as the center and select all the atoms in a 8Å radius. The features of these atom tokens are average pooled, passed to the classifier, before being normalized into the 20 amino acid likelihoods. The classifier consist of a Linear-ReLU-Linear block, with 128 and 20 neurons at these linear layers.

### ΔΔ*G* Predictor

Our ΔΔ*G* predictor contains two parts, a backbone which encodes the masked local environment and a regression head. For the purpose of this discussion, we primarily focus on the architecture of the regression head, as the pre-trained weights from the self-supervised learning task are loaded into the backbone. We get one masked local environment together with two amino acid types (one is the wild-type and the other is the mutation), and target at predicting the Δ*G* changes.

We first extract useful atomic features from the backbone model. Given a masked local environment, we pass it through the backbone and get the output *h ∈* R*^N×H^* where *H* denotes the hidden dimension. Given the wild-type and mutation amino acid type, we extract the corresponding amino acid embedding *e_wt_ ∈* R^1^*^×H^* and *e_mut_ ∈* R^1^*^×H^* in the final linear layer of the backbone. We then apply Concat(*h, e_wt_*) and Concat(*h, e_mut_*) to get two hidden representations for wild-type local environments and mutated local environment, respectively.

Pass Concat(*h, e_wt_*) and Concat(*h, e_mut_*) through the attention blocks, we extract the *e_wt_*and *e_mut_*in the final layer, substract them, apply a linear layer and output the predicted ΔΔ*G*.

### Training Configuration

During training, we shuffle the training data and use Huber loss with *δ* = 1 as the training loss. When training on large-scale datasets, *e.g.*, cDNA120K, we end-to-end tune all the parameters, with learning rate 5 *×* 10*^−^*^5^ for regression head parameters and learning rate 2 *×* 10*^−^*^5^ for backbone parameters. We apply optimizer AdamW with batch size 960 (batch size 240 with accumulation step 4), weight decay 0.1, optimizer EMA with *η* = 0.99, number of iterations 750. To finetune the CDNA-trained model on the small datasets (*e.g.*, C2878), we freeze the backbone parameters and use AdamW with learning rate 5 *×* 10*^−^*^7^, batch size 1, 024, weight decay 0.1, number of iterations 500. Train from scratch on the small datasets (*e.g.*, C2878), we freeze the backbone parameters, and set optimizer AdamW with 5 *×* 10*^−^*^5^, batch size 1, 024, weight decay 0.1, and number of iterations 500.

### B.2 Experiment Settings

#### Pretraining Dataset

We pre-train our graph-transformer backbone using the same procedure as MutComputeX [50]. Briefly, this dataset consist of a 90:10 split of 2,569,256 microenvironemnts sampled from 22,759 protein sequences clustered at 50% sequence similarity from PDB structures with up to 3Åresolution.

#### Training Set

We clean up a new dataset cDNA120K [46], which contains 144K labeled free energy data for small proteins. We first train on this dataset and fine-tune it on C2878.

#### Test Set

To fully test different models, we collect and clean related datasets in the literature and make our new version test set. To get more comparisons, we report the numbers on all the literature datasets here. In summary, we test our model on S-sym, P53, Myoglobin and S669. S-sym, P53, and Myoglobin datasets are proposed in ThermoNet [28] and are widely used by the follow-up works as the testing benchmarks. S669 includes curated data dissimilar at 25% sequence identity to S2648 training data. Additionally, we create several more domains from the original test data. ➀ Swap the mutation and wild-type amino acids, and this new test domain is marked as TR. ➁ Once we have multiple single-point mutations in the same position, we randomly pick two amino acids as the wild-type and the mutation. We mark this test domain as TP.

#### Evaluation Metrics

We evaluate regression and classification metrics. First, for a fair comparison, we report Pearson correlation and RMSE for our regressor. Furthermore, in practice, instead of RMSE, we have more interest in the performance on stabilizing mutations. Therefore, we report the precision and recall for the stabilizing mutations and the AUROC values for the binary classification problems.

### C Overview of Deep Learning for Stability Prediction

The lack of available stability data has prevented deep learning frameworks from significantly accelerating the field and the community still primarily relies on knowledge-based methods, such as Rosetta [32] and FoldX [33], or shallow machine learning approaches [34, 35, 36, 37, 38, 39, 40, 41, 42]. Nonetheless, several deep learning frameworks for stability predictions have been published and they can be broadly categorized by their input type: sequence or structure. The current structure frameworks include DeepDDG [29], ThermoNet [28], ProSGNN [30], and AC-DNN [27] and sequence frameworks include Prostata [26] and HotProteins [73]. While both structure and sequence frameworks are competitive with current state-of-the-art methods, the two types have their own benefits and limitations. The structure frameworks capture atomic interactions within a protein and with other biomolecules, such as ligands, nucleotides, and other proteins. Furthermore, the richness of atomic representations enables generalization while training with only 10^3^-10^4^ protein structures. However, all existing frameworks require an additional mutant structure to make a stability prediction. This adds a significant computational hurdle that scales linearly with the number of inferences. For example, to conduct a computational deep mutational scan (DMS) of a 300 amino acid protein, one must computationally generate 5700 mutant structures via AlphaFold2 or Rosetta[32]. For sequence-based frameworks their primary benefit is that they required little data engineering and have billions of primary sequences readily available for training. This gives them a low barrier to entry and makes them popular amongst the machine learning community, resulting in an explosion of transformer-based large language models (LLMs) across the protein community[63, 64, 25, 70, 26, 68, 65, 69, 74, 75]. There are two primary limitations to sequence models. First, LLMs require extensive training on large sequence databases (UniRef50 *∼*46M) to learn meaningful representations for downstream applications. Training these models is very computational expensive and only affordable to well-funded academics and technology companies [63, 64, 25, 68, 69, 75]. Second, the self-supervised pre-training of transformer models constrains the training space to single protein sequences. As protein engineers, we intuitively understand that proteins are more than their primary sequence; proteins often form complexes, undergo post-translational modifications, and interact with cofactors, DNA, RNA, and oligosaccharides. This artificial domain shift limits their generalization in downstream task and has already been empirically demonstrated when predicting stability changes at protein-protein and protein-ligand interfaces [26].

## D Data and Metric issues in Computational Stability Prediction

### D.1 Data

Systematic analysis of computational stability predictors published over the last 15 years has identified several important shortcomings hindering the field from taking advantage of deep learning algorithms. These shortcomings are present in every facet of the data pipeline: data scarcity, data bias, data leakage, poor metrics for evaluating performance, and computational cost. Here is a high-level, non-exhaustive overview of the most pressing issues. The limited amount of experimental data is heavily biased in several ways: 1) dominated by destabilizing mutations, resulting in only *∼*20% of point mutations predicted to stabilize indeed increase stability [14, 31, 13]; 2) stabilizing mutations are narrowly concentrated between −2 to 0 kcal/mol, causing models to overfit to this distribution and underestimate and not generalize to strongly stabilizing mutations [14, 31, 13]; 3) stabilizing mutations are enriched with surface hydrophobic mutations, causing stabilizing predictions to often unintentionally lower protein solubility [14, 31, 13] 4) of the 380 types of amino acid substitutions *∼*20-70% have no data in the common training and validation sets and the ones sampled are significantly biased toward mutations “from” or “to” alanine and valine [31]. Due to the hierarchical nature of proteins, splitting experimental data into training and validation sets can be done at the mutation, position, protein, and sequence cluster levels. All but sequence cluster result in data leakage, thus, most published computational tools report inflated evaluation metrics and display poor generalization.

### D.2 Metrics

The common metrics used to assess the performance of computational stability predictors (Pearson correlation, classification accuracy, and error) are not appropriate for the goal of accurately identifying stabilizing mutations and tend to be misleading [14, 31, 13]. The metrics naively formulate the problem as a regression or classification task and are not tailored to the actual goal: accurately predicting stabilizing mutations. This misalignment has resulted in metrics that do not prioritize improvement in accurately predicting stabilizing mutations and their game-ability has been demonstrated in a study that consisted of 21 established computational tools [31]. Here, the authors demonstrate how Spearman and Matthew correlation can correctly identify deficiencies in an overly simplified model while the original metrics couldn’t. We argue that these metrics are still insufficient, and metrics specifically tailored toward accurately identifying strong stabilizing mutations are still required.

